# Engineered human mesenchymal stem cells as new vaccine platform for COVID-19

**DOI:** 10.1101/2020.06.20.163030

**Authors:** Junhua Liu, Huping Jiao, Xiushan Yin

## Abstract

Recently, there are several routes for COVID-19 vaccine research, yet their weaknesses lie in low efficiency, tolerability, immune adaptability and safety. We describe a new approach to COVID-19 based on engineered human mesenchymal stem cells(hu-MSC), which is like a small protein antigen response device, but will be gradually cleared and degraded by body’s immune system among recognization process. The antibody response results show that this is effective and fast. Furthermore, after several antibody response tests, we obtained an injection of a set of cocktail-like modified human mesenchymal stem cell line. This strategy opened a new avenue for vaccine design against COVID-19.

## Introduction

The pandemic of COVID-19 has spread around the world and vaccines become the main means for fighting of COVID-19^[1-3]^. At present, hundreds of vaccines are under the pipeline including traditional attenuated/inactivated vaccine, nucleic acid-based vaccine (DNA and RNA^[4]^), and protein vaccine^[5-7]^. Each of these vaccines has their own advantages and disadvantages^[8-11]^, but the vast majority of vaccines need to be repeatedly injected to immunize the body, during the response period some causes the body’s uncomfortable symptoms. Some lack the modification happened in vivo during virus replication. Until today it’s not clear whether the vaccines developed so far will be successful or not, since none of the vaccines had been developed for other human coronavirus. Therefore, the development of a safe and effective vaccine may be a key to control the COVID outbreak and future novel coronavirus^[1, 10]^. Here we demonstrated the modified MSCs expressing SARS-CoV-2 proteins can be used as a novel vaccine platform.

## Results and discussion

Firstly, we took advantage of allogenic mesenchymal stem cells (MSCs) that are widely used in transplantation-based cell therapy^[12]^. Human umbilical cord derived MSCs were isolated and evaluated with MSCs typical features characterized by FACS and functional assay (data not shown). Then primary MSCs at passage four were transfected with plasmid expressing nucleocapsid (N) protein, the structural protein showed dominant and long-lasting immune response reported for SARS-CoV (outbreak in 2003)^[13]^. Expression was evaluated and transfection efficiency was approximately 20% on average after 48 hours of transfection. The MSC-SARS-CoV-2-N cells were used for mice injection. In total one million modified MSCs were injected per mice (C57BL/6) via intramuscular (IM) or subcutaneous (SC) routes, respectively, to allow vaccine antigens being produced in vivo (Figure 1).

**Figure 1.**
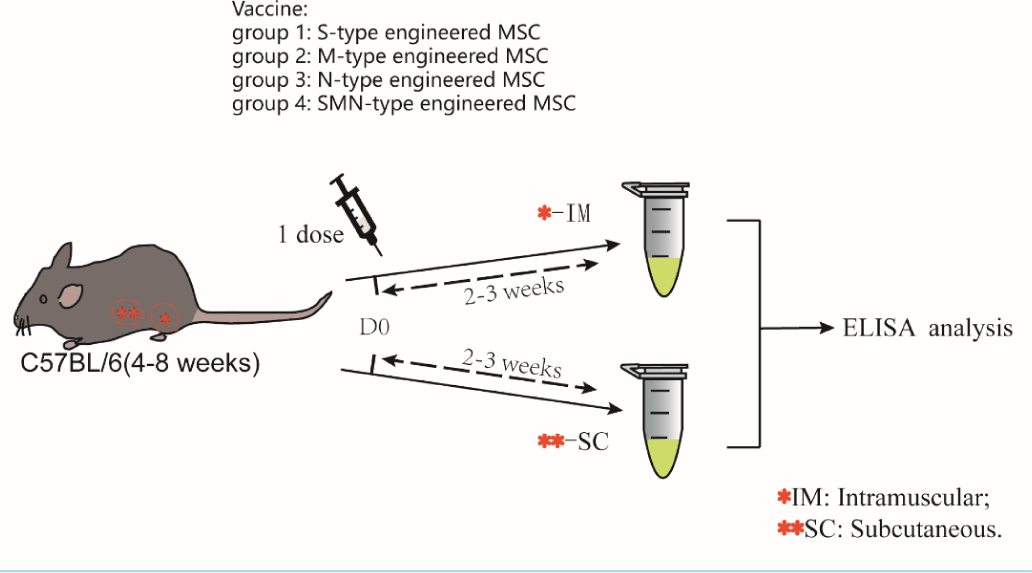
A schematic diagram of the one-dose MSCs injection. Four groups of mice were given a single dose of intramuscular injection with the several type engineered MSC. Similarly, the other four groups of mice were given another subcutaneous injection for the same experiment. After two weeks later, blood sample was collected for ELISA.

The results showed that after 20 days, almost all the immunized mice had antibody production in serum (7/8), and at least 50% of the mice showed strong positive antibody expression followed by IM or SC injections with one dose injection (Figure 2). The response for IM or SC injection was comparably judged by ELISA and with similar efficiency. Control mice injected with empty vector did not show anti-N response, and the ELISA signal is comparable to background with none-injected mice (Figure 2). We did not observe any severer symptoms from the injected mice after any injection. And the body weight of mice during this study were remained stable or increased. Successful antibody response might due to the adaptive immunity promotion effect of MSCs described before^[14]^. We then co-transfected the S, N and M, the structure proteins of SARS-CoV-2, and observed elevated antibody response for N protein again. Due to lack of good antibodies used for ELISA for S and M protein, we were not able to evaluate the effect of these two antibody responses. And potential more elevated signal should be obtained by multiple injections.

**Figure 2.**
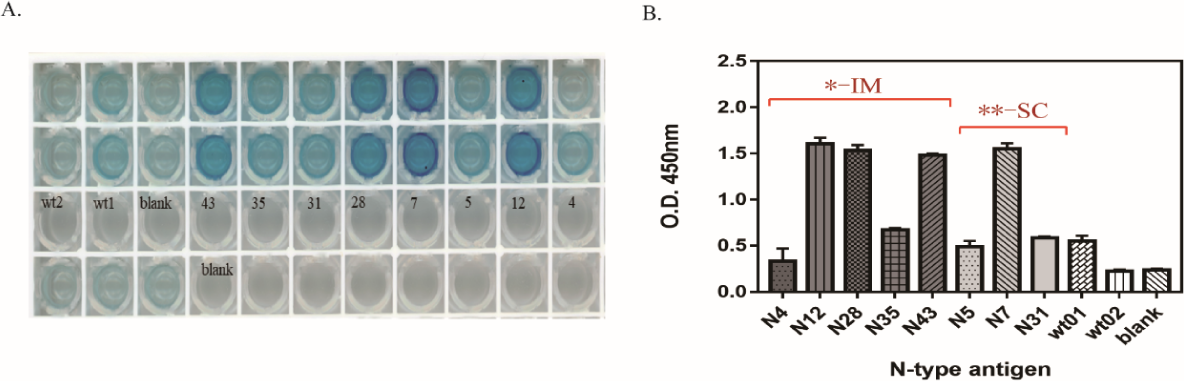
The serology antibody detection after engineered hu-MSC immunization. The figure 3A shows the color development of serum of mice immunized with N-type MSC vaccine in ELISA. The shade of color represents the content of antibody. Among them, there is a slight background value of blank or negative control, which may be due to low purity of antigen protein. the figure 3B shows that the absorbance values at 450nm after immunization with N-type MSC vaccine. SC: subcutaneous(the numbers of mice were N4, N12, N28, N35, N43); IM: intramuscular(the numbers of mice were N5, N7, N31).

One potential risk for cell-based vaccine is that residual cells in the body might cause uncontrolled proliferation and malignancy. PCR for N gene sequence and human gene sequence were performed in blood and tissues from injected region, and no positive signal was detected, indicating that the MSCs were cleared by immune system after 20 days of injection (data not shown).

The research and development route based on stem cells proposed in this report can realize the presentation of antigen proteins of multiple targets by virtue of the feature of presenting and secreting part of the novel coronavirus antigen protein. Most importantly, some of the novel coronavirus proteins mediated and secreted by stem cells will be modified, like glycosylation^[15]^ and acetylation, mimicking the virus protein modification in vivo, which can well simulate response increase the success rate of one-time immunity^[16]^. It was reported that mice strains maintained in the laboratory are not susceptible to SARS-CoV-2 infection^[17]^, due to the variation of ACE2 receptor for virus entry^[18, 19]^. Later other model animals^[20]^ like dogs and monkeys are needed to further test the efficiency and performance of stem cell vaccine. To our knowledge, this is the first report on a novel vaccine against COVID-19 based on stem cell as a platform. Combined with the advantages of mesenchymal stem cells in the treatment of diseases ^[21]^, we believe that such stem cells or cell-based vaccines will be a promising strategy to control COVID-19 and other coronavirus with unknown potential risk^[22]^, and it is also a promising model for some pandemic disease.

## Materials and Methods

### Animal

All mice in our experiment were provided by Changsheng Animal Research Institute in Liaoning Province, and Male and female C57BL/6 mice (4-8 weeks old) were used for immune injection. Animal works were approved by ethical committee of Shenyang University of Chemical Technology.

### Hu-MSC cell isolation

The fresh umbilical cord tissue was cut into some pieces and isolated enzymatically in an incubation solution of 10mL Trypsin-EDTA (Cat: 25200-072, Gibco) for 40 min at 37°C. The digestion homogenate was stopped by 10mL DMEM/F12(Cat:11039-021, Gibco) with 10% FBS (10099141C, Gibco) and then slightly decentralized 30-40 times through 1-mL pipette tips. Finally, all cell solution was filtered by 100 μm cell strainer (Cat: 352360, Corning FALCON) into a new centrifugal tube, and the P0 hu-MSC was isolated.

### Generation of vectored antigen protein by eukaryotic expressing system

Refferring to the SARS-COV-2 virus genome sequence from Wuhan-Hu-1 annotated in Genebank (locus:NC_045512.2 or MN908947)^[23]^, we synthesized the spike gene, membrane gene and nucleocapsid gene in gene synthesis company (GENEWIZ, china). In terms of S type and N type plasmids, these long gene sequences containing enzyme sites firstly was amplified by common PCR by Q5 high-fidelity PCR kit (Cat: #M0491, NEB), using primer F and R pairs of S or N. Subsequencely, these amplication products were added to the linear eukaryotic expression vector pcDNA3.1-flag, by the way of the restrictive enzyme digestion-ligation. However, since the M gene was limited by the above selecting restriction sites, such as NotI and Xbal, we chaged the homologous recombination method. A third M type primers were used to amplify the M sequence carrying the linear vector’s arms. The PCR product was purified and directly recombined by the EasyGeno recombinant clone kits (Cat:VI201, Tiangen). After conventional transformation and monoclonal screening of positive bacteria, we obtained the correct plasmids construction. Finally, the plasmid construct were purified by OMEGA Free-endotoxin extraction kits (Cat: D6950, OMEGA), and eluted in the RNase-free water. All primers were as follows:

S-FP: 5’-GGGCGGCCGCCCTTTGTTTTTCTTGTTTTATTGC-3’

S-RP: 5’-CCCTCTAGAGCTTATGTGTAATGTAATTTGACTC-3’

N-FP: 5’-GGGCGGCCGCCCTCTGATAATGGACCCCA-3’;

N-RP: 5’-CCCTCTAGATTAGGCCTGAGTTGAGTC-3’;

M-FP: 5’-GACGACAAGGGGCGGCCGCAAGCAGATTCCAACGGTACTAT-3’;

M-RP: 5’-TTTAAACGGGCCCTCTAGATTACTGTACAAGCAAAGCAAT-3’.

### Cell culture, Plasmid transfection and MSC-vaccine preparation

All human mesenchymal stem cells were cultured in free serum medium (Cat: SC2013-G-A, AM-V serum free medium) supplement with related cytokines and 1% pennicillin-streptomycin (Cat:MA0110, Meilunbio). These cells were refreshed medium or passaged every three days. In our experiments, we usually choose the fourth or sixth passage of hu-MSC for transfection. Due to the difficulty of transfection, we adopted the method of electro-transfection to deliver the plasmids into hu-MSC. After transfection, the MSC plates were incubated at 37°C with 5% CO2.

When the engineered hu-MSC cultured until 48 hours post-transfection, the adherent MSC were digested using 0.25% trypsin-EDTA and cell pellets were harvested. The modified hu-MSCs were resuspended with PBS and counted to make dilutions to form proper concentration.

### Engineered hu-MSC injection and in vivo immunization

An equal amount of male and female C57BL/6 mice, aged from six to eight weeks, were taken. Then, the mice were randomly divided into two groups, according to the two injection ways (intravenous and subculaneous). For example, the right hind leg of the mice was sterilized by 75% alcohol and injected 1 million modified MSC through 1 ml injector for intramuscular, and choosing the side abdomen or neck locus were for subcutanous injection (SI). Briefly, no matter what kinds of injection, we would use 1 million engineered hu-MSC for vaccination.

### FACS analysis

To evaluate the feasibility of the transfection condition, we delivered a plasmid carrying GFP as a reporter to hu-MSC though electro-transfection, and the transfection efficiency was measured by flow cytometry (FACS). Generally, we added 0.25% Trypsin-EDTA about 500 µl to one well of six-well plate, digested and separated, harvested cell pellet. Then, the cells were resuspended with appropriate PBS buffer, and filtered through a 100 µm strainer for detection on flow cytometer.

### ELISA analysis for antibody

Blood sample were taken from the mice for serological test, after 20 days of vaccination. Meanwhile, we randomly selected 5-8 wild mice without any treatment from the same batch of mice before injection, and their serum were performed as negative controls. We assessed the ability of mice to produce antibodies against nucleocapsid protein by ELISA analysis. ELISA 96 well plates were coated with specific antigen protein (the dilution ratio: 1/20-30), which manufactured by Beijing sino biological(cat:40589-v08B1). Then, sample diluent 100 µl and every sample serum 5 µl were added to each well of 96 well plate, and incubated at 37°C for 60 min. After 1 hour incubation, solution was discarded and the plate was washed by PBS for 5 times. The secondary HRP-labeled Goat Anti-mouse IgG(H+L) (Cat:A0216, Beyotime) at the dilution ratio of 1 to 10000 was added to each well and incubated at 37°C for 30min. The same washing was performed for 5 times. After TMB (Cat:ST746, Beyotime) was added for color development for 15min, absorbance was detected under 450nm condition via Multiscan Spectrum when added with stopping buffer.

## Statistical analysis

For gene sequence, we successively adopt SnapGene to analysis the sequencing results. Then, we used Graphpad Prism V5.0 analysis for the group values to draw the graph, and from which experimental data were presented as the mean ± SD and analyzed by the unpaired Student’s t test.(P<0.05 was considered statistically significant).

